# What can genomics tell us about the success of enhancement programs in anadromous Chinook salmon? A comparative analysis across four generations

**DOI:** 10.1101/087973

**Authors:** Charles D. Waters, Jeffrey J. Hard, Marine S.O. Brieuc, David E. Fast, Kenneth I. Warheit, Robin S. Waples, Curtis M. Knudsen, William J. Bosch, Kerry A. Naish

## Abstract

Population enhancement through the release of cultured organisms can be an important tool for marine restoration. However, there has been considerable debate about whether releases effectively contribute to conservation and harvest objectives, and whether cultured organisms impact the fitness of wild populations. Pacific salmonid hatcheries on the West Coast of North America represent one of the largest enhancement programs in the world. Molecular-based pedigree studies on one or two generations have contributed to our understanding of the fitness of hatchery-reared individuals relative to wild individuals, and tend to show that hatchery fish have lower reproductive success. However, interpreting the significance of these results can be challenging because the long-term genetic and ecological effects of releases on supplemented populations are unknown. Further, pedigree studies have been opportunistic, rather than hypothesis driven, and have not provided information on “best case” management scenarios. Here, we present a comparative, experimental approach based on genome-wide surveys of changes in diversity in two hatchery lines founded from the same population. We demonstrate that gene flow with wild individuals can reduce divergence from the wild source population over four generations. We also report evidence for consistent genetic changes in a closed hatchery population that can be explained by both genetic drift and domestication selection. The results of this study suggest that genetic risks can be minimized over at least four generations with appropriate actions, and provide empirical support for a decision-making framework that is relevant to the management of hatchery populations.

## Introduction

Enhancement, the release of cultured organisms to increase population abundance, is an important fishery management tool (Lorenzen *et al*. 2010). But genetic risks associated with artificial propagation are well known and may compromise the wild populations that enhancement is intended to support (Naish *et al*. 2008; Laikre *et al*. 2010). Supportive breeding programs are a form of enhancement used extensively in the management of Pacific salmon in North America. Such programs aim to increase population sizes by rearing a fraction of juveniles in captivity and then releasing them into the natural environment along with their wild-born conspecifics (Ryman & Laikre 1991). Concerted efforts have been directed at mitigating the effects of domestication selection, genetic drift and inbreeding (Mobrand *et al*. 2005) associated with these programs, because in many cases populations have not recovered and cannot support sustainable fisheries (Naish *et al*. 2008; Beamish *et al*. 2010; Scheuerell *et al*. 2015). Practical recommendations to mitigate genetic risks have focused on theoretical models that examine the influence of gene flow in reducing divergence between cultured and wild populations (Duschene & Bernatchez 2002; Ford 2002; Baskett & Waples 2013). Specifically, the intentional use of natural-origin broodstock in the creation of the hatchery population in each generation may reduce risks, especially when gene flow from the hatchery to the wild population is limited (Mobrand *et al*. 2005). Such “managed gene flow” has seen widespread adoption in the Pacific Northwest of the USA (Paquet *et al*. 2011), but few practical examples on their efficacy exist.

An ideal way to test whether managed gene flow is effective at reducing genetic divergence between hatchery and wild populations is to empirically compare cultured populations with and without gene flow. Such a comparison would provide results on the range of possible outcomes of these management approaches, and would be especially informative if conducted longitudinally. The use of population genomic approaches provides a way to survey temporal changes in genetic divergence, to measure the rate of change with each generation since founding, and to identify factors driving divergence. We previously conducted such a study in two populations of Chinook salmon (*Oncorhynchus tshawytscha)* released from a hatchery on a tributary of the Columbia River (Waters *et al*. 2015). Both hatchery populations were founded from the same source population; however, one population remained integrated with the wild and used only wild-born broodstock in each generation, while the second hatchery population was maintained separately and received no gene flow from the wild. Our earlier results over three generations revealed little change in the integrated line compared to the founding population. Most of the genetic divergence in the segregated line could be attributed to genetic drift, but there was also evidence for directional selection at specific locations in the genome. However, it is unclear over how many generations managed gene flow may be effective at mitigating genetic risks, because processes occurring in the wild could mitigate or exacerbate the effects over time (Ford 2002; Baskett & Waples 2013). Here we aimed to test whether the use of natural-origin broodstock was effective at reducing divergence over several generations by extending our earlier study for an additional fourth generation.

## Materials and Methods

A spring Chinook salmon hatchery program was initiated in 1997 at the Cle Elum Supplementation and Research Facility (CESRF, Fig. 1) to supplement the declining upper Yakima River Chinook salmon population while minimizing possible genetic and ecological risks associated with supportive breeding. Local, wild adults were collected for broodstock from 1997 to 2002 as they passed the Roza Dam Adult Monitoring Facility (Roza Dam, Fig. 1). Adults were then transferred to CESRF and held until spawning. Eggs and juveniles were reared in the hatchery for approximately 18 months before they began their migration to the ocean. Adult hatchery fish first returned to the Yakima River in 2001 and were allowed to spawn naturally. In 2002, both wild and hatchery-origin adults were spawned at CESRF to create two contrasting hatchery lines. The integrated (INT) line is derived only from wild or natural-origin adults, and all fish from this line are allowed to spawn naturally. Here, natural-origin fish are those that were born in the river but may have some hatchery ancestry. The segregated (SEG) line, however, uses only hatchery-origin broodstock, and no fish can spawn in the river.

**Figure 1.**
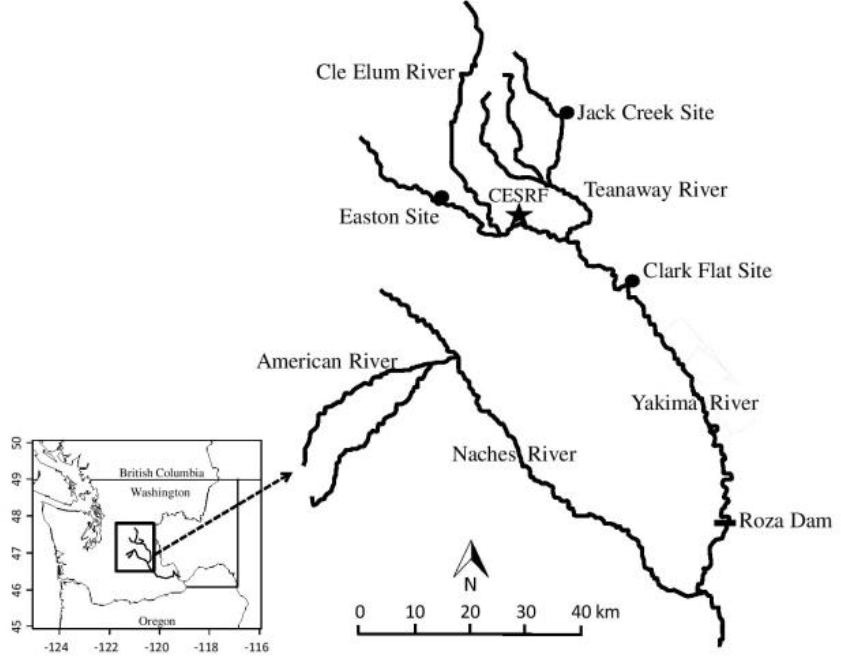
From Waters *et al*. (2015). Map of the Yakima River system. The upper Yakima Chinook salmon population is the target of the Cle Elum Supplementation and Research Facility (CESRF). All returning adults are sampled at Roza Dam and allowed to spawn naturally (natural origin and integrated line fish) or are removed from the system (all segregated line fish). Spawning and rearing for the hatchery lines occurs at CESRF. Prior to outmigration in spring, juveniles are transferred to the Easton, Jack Creek, and Clark Flat acclimation sites, where they are held for approximately two months before volitional release.

Tissues for DNA were sampled from adults of both hatchery lines in 2014 during spawning at CESRF and stored in 100% ethanol. These adults represent the fourth (F_4_) generation of each line. DNA was extracted using DNeasy Blood & Tissue kits (Qiagen, Valencia, CA, USA) following the animal tissue protocol. Restriction site-associated (RAD) libraries (Baird *et al*. 2008) were prepared using the restriction enzyme *SbfI* and sequenced on the Illumina HiSeq 2000 platform with 36 individuals per lane. All raw RAD sequences from the F_4_ generation were combined with raw data from our previous comparative analysis (P_1_ founders and F_1_-F_3_ generations, Waters *et al*. 2015). Filtering and genotyping were performed following Waters *et al*. (2015), with two additional steps to improve data quality. First, loci were removed if more than 50% of individuals in any population were not genotyped. Then, loci were removed if they did not meet Hardy-Weinberg equilibrium conditions (*q*-value < 0.05) in more than one population, as determined by the Monte Carlo procedure with 1x10^5^ permutations in the R-package *adegenet* (v. 1.3-9, Jombart 2008). g-values were computed using the R-package *Q-value* (v. 1.28.0, Storey 2002).

As the aim of the present study was to extend previous comparisons between the integrated and segregated hatchery lines by another generation, genetic change was evaluated using the same methods described in Waters *et al*. (2015). Population-level genetic change between each generation of the hatchery lines was evaluated using measures of *F_ST_*, computed in *Genepop* (v. 4.1, Raymond & Rousset 1995), and a discriminant analysis of principal components (DAPC), conducted in the R-package *adegenet.* The relative effect of genetic drift within each hatchery line was determined using estimates of effective numbers of breeders, N_b_. Temporal and linkage disequilibrium (LD) estimates of Nb were computed with N_E_ Estimator (v. 2.01, Do *et al*. 2014) using only four-year-old adults, which represented a single cohort of individuals. Steps taken to reduce potential bias in Nb estimates due to selection, overlapping generations, and fluctuating population size were identical to those of Waters *et al*.(2015). Lastly, loci and genomic regions exhibiting signals of diversifying selection in the hatchery lines were identified using three independent tests: *F_TEMP_* (Therkildsen *et al*. 2013), *Bayescan* (Foll & Gaggiotti 2008), and a sliding-window approach (Brieuc *et al*. 2015; Waters *et al*. 2015). We focused on loci and regions that were identified by multiple tests and were divergent across multiple generations.

## Results

Tissues from 72 individuals (36 from each line) were sequenced from the F_4_ generation. The raw data was combined with RAD sequences from the previous generations and filtered, yielding 9266 bi-allelic RAD loci with minor allele frequencies >0.05 in at least one population and less than 50% missing genotypes within each population. A total of 465 individuals from the five generations were genotyped at >50% of these loci and retained for analyses (Tables S1, S2). Tests of HWE identified 158 loci that significantly deviated from expectations in more than one population. Following removal of these loci, the final data set comprised 9108 loci (Table S1), including 4214 loci that aligned to the Chinook salmon linkage map (Brieuc *et al*. 2014). Population-level divergence of the two hatchery lines followed previously documented trends (Waters *et al*. 2015). Values of pairwise *F_ST_* between the lines and the P_1_ founders was approximately four times higher in the F_4_ SEG population (*F_ST_* =0.0125, P < 0.001, Table S3) than in the F_4_ INT population (*F_ST_* =0.0033, P < 0.001). Divergence between the two hatchery lines in the F_4_ generation also continued to increase (*F_ST_* =0.0126, P < 0.001). Patterns of genetic change were further supported by a discriminant analysis of principal components, conducted on the first 63 PCs as recommended by the *optim.a.score* function in *adegenet.* The segregated line diverged from the P_1_ founders and integrated line over time along the first discriminant function; this axis explained 59.2% of the retained variation (Fig. 2). Genetic change between the later generations of the integrated line and the P_1_ founders was evident along the second discriminant function, which explained 16.6% of the retained variation.

**Figure 2.**
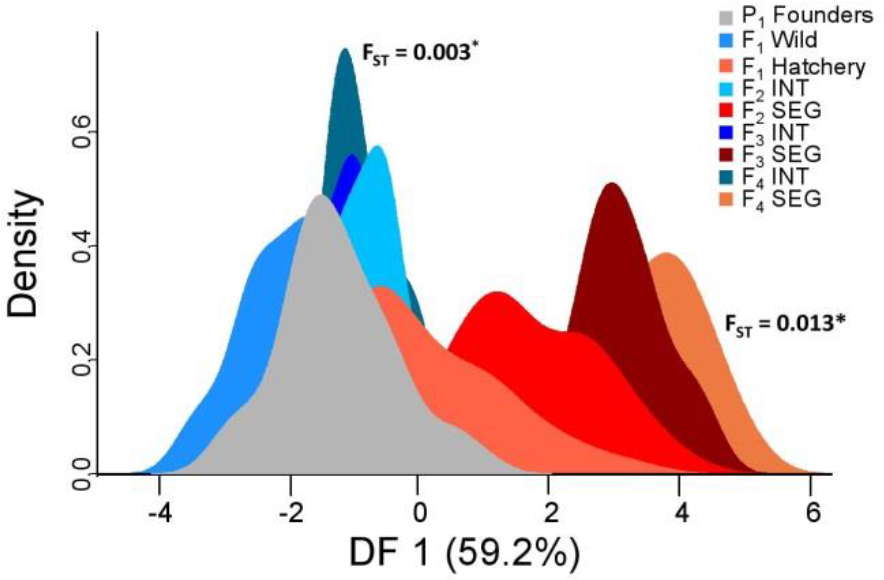
Density plot of individuals along the first discriminant function from the discriminant analysis of principal components (DAPC) for the wild founders (P_1_ Founders, black) and four generations of the integrated (INT, blue colors) and segregated (SEG, red colors) hatchery lines. Pairwise F_*ST*_ values for the F_4_ generation compared to the P_1_ founders are shown for each hatchery line.

Bias-adjusted LD and temporal estimates of effective number of breeders, N_b_, supported earlier results and suggested that the relative effect of genetic drift was much greater in the segregated line than in the integrated line. The LD estimate of N_b_ in the F_4_ INT population was nearly eight times higher than that obtained in the F_4_ SEG population (Fig. 3; Tables S4a, S5). The temporal N_b_ estimates, which applied to the entire sampling period of 1998-2014, were also markedly different between the two lines (Fig. 3; Table S4b). While the average broodstock size of the integrated line (363 ± 15) exceeded that of the segregated line (85 ± 15), the difference was not sufficient to explain the higher N_b_ of the integrated line (Table S6).

**Figure 3.**
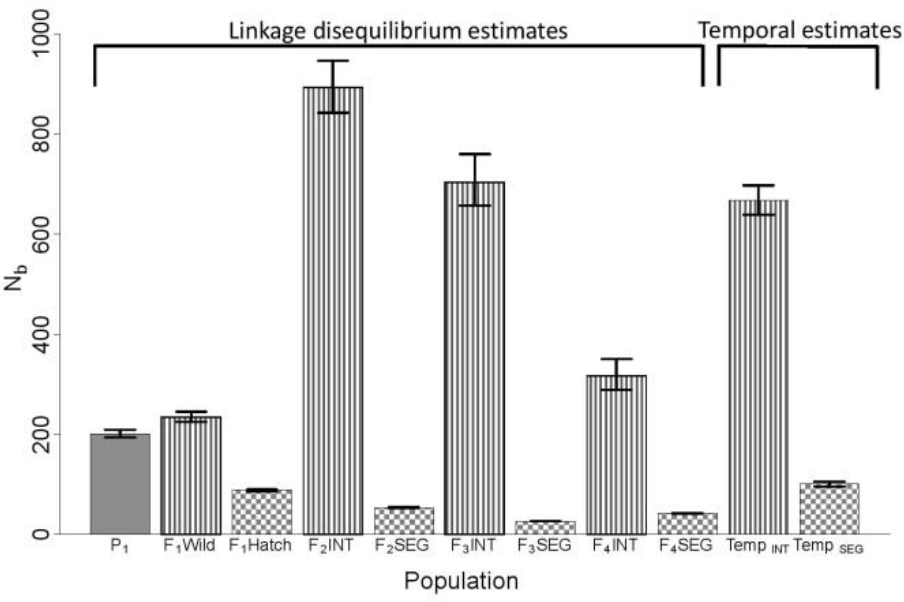
Estimates of effective number of breeders, N_b_, and 95% confidence intervals produced by the linkage disequilibrium (LD) and temporal methods. The LD method enables estimation of N_b_ for every generation, while a single estimate for the sampling period is produced by the temporal method. LD estimates are adjusted for physical linkage and other potential biases as described in Waters *et al*. (2015).

In addition, the Nb estimates for the F_4_ generation revealed a result that was not apparent from the census data alone. The ratio of effective number of breeders to census size (N_b_/ N_census_) declined from 0.21 (95% CI: 0.20-0.23) in the F_3_ INT sample to 0.04 (95% CI: 0.03-0.04) in the F_4_ INT sample, despite the fact that the census sizes increased from 3364 to 8374 adults. This result is important because the N_b_/N_census_ ratio provides a metric for understanding factors which cause deviations from N_b_=N_census_ (e.g. variance in reproductive success) and affect genetic variation over time.

Three independent tests identified loci and genomic regions that exhibited signals of diversifying selection. The *F_TEMP_* method identified 78 loci that exceeded neutral expectations in the integrated line and 198 in the segregated line (Table S7). Thirty-five loci were outliers in both hatchery lines. *Bayescan,* conducted using all populations combined, identified 120 loci putatively under diversifying selection (Table S7). There was considerable overlap between *Bayescan* and F_TEMP_, as 48 and 72 *Bayescan* outliers were also identified by *F_TEMP_* in the integrated and segregated lines, respectively. Genomic regions that exhibited significantly elevated levels of divergence compared to the P1 founders were identified in both hatchery lines by sliding window analyses (Table S8). However, divergence in the segregated line was more consistent across the F_1_, F_2_, F_3_, and F_4_ generations than in the integrated line. For example, seven regions were significantly elevated in at least three generations of the segregated line while none were observed in the integrated line (e.g. Fig. 4). Five of these regions also contained outlier loci identified by *F_TEMP_* and *Bayescan* (Fig. 4), providing further support that selection - likely due to continued exposure to the hatchery environment - has also contributed to the higher levels of divergence observed in the segregated line. Previous work has identified genes in such regions of overlap that may be targeted by selection in captivity (Waters *et al*. 2015).

**Figure 4.**
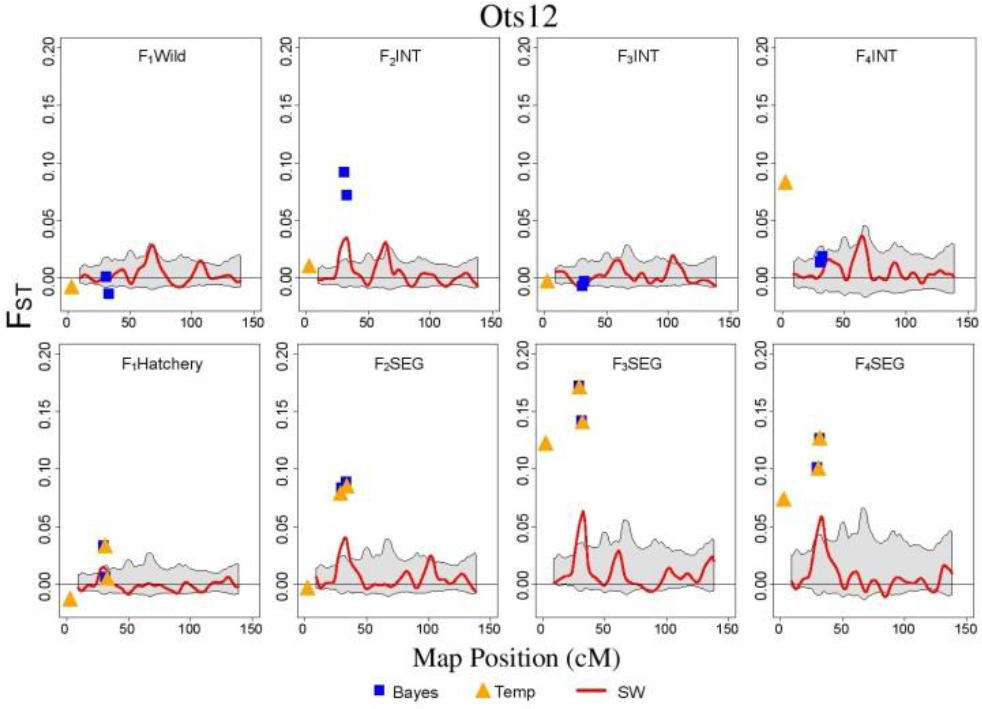
Loci and regions of the genome showing signatures of adaptive divergence, based on pairwise *F_ST_* compared to the P1 founders, on chromosome Ots12 for the integrated (top panel) and segregated (bottom panel) hatchery lines through the F_1_, F_2_, F_3_, and F_4_ generations. Blue squares are loci that were identified as outliers with *Bayescan*and orange triangles are outliers identified by *F_TEMP_.* The red line represents the kernel smoothed moving average of *F_ST_* and the grey shaded area is the 95% confidence interval.

## Discussion

Here, we have demonstrated the utility of genomic-based methods to test alternative management approaches for population enhancement and to monitor fine scale genetic changes in populations over several generations. Many theoretical studies have indicated that ongoing gene flow between hatchery and wild fish may ultimately compromise the fitness of the natural population (Ford 2002; Baskett & Waples 2013). However, the degree to which the natural population is affected depends on many factors that likely fluctuate over time, such as selection intensity, the proportion of wild-origin individuals on the spawning grounds, reproductive rate in the hatchery and wild, and carrying capacity of the natural system. Thus, our multigenerational findings extend complementary studies that evaluate reproductive success in single populations over one or two generations (Christie *et al*. 2014). The results from the fourth hatchery generation largely supported observations from a previously published longitudinal study (Waters *et al*. 2015). Little genetic change occurred in the integrated hatchery line, which frequently exchanged migrants with the founding wild population. In contrast, consistent temporal trends in divergence were documented in the segregated hatchery line, which is maintained as a closed population. Such consistency was observed on the population-level and at specific genomic regions, despite the fact that environmental change likely occurred during the sixteen years over which this study was conducted. This result might be explained by domestication selection imposed by the relatively uniform hatchery environment on the segregated hatchery population. Genomic regions exhibiting potential signals of domestication selection can be further examined to identify candidate genes (e.g. Waters *et al*. 2015) and mechanisms underlying genetic adaptation to captivity, and to inform management practices to possibly reduce this risk.

Notably, extending the earlier study by another generation also revealed fluctuations in N_b_/ N_census_ that would otherwise have been missed. We previously reported Nb/Ncensus ratios of 0.11 (95% CI: 0.10-0.12) and 0.21 (95% CI: 0.20-0.23) for the F_2_ and F_3_ INT samples, respectively, which reflected the first two generations of naturally-spawning adults that included hatchery fish from the integrated line. These estimates show a positive trend in Nb/Ncensus, which, if taken alone, could possibly be attributed to successful supplementation efforts. However, it is important to acknowledge temporal variability, and the decline of N_b_/N_census_ to 0.04 in the F_4_ INT sample may have two explanations. The first is that the results may indicate the influence of the Ryman-Laikre effect (Ryman & Laikre 1991), where supportive breeding reduces the effective size of a wild population. Alternatively, temporal fluctuations in N_b_/N_census_ could simply reflect changes in the natural environment that influence demographic factors; the observed ratios of 0.04-0.21 are within the range of those documented in natural populations of many species (including salmonids; Frankham 1995; Naish *et al*.2013). It is impossible to identify the true source(s) of the observed fluctuations, particularly since there is no wild control population for comparison. Nevertheless, our results emphasize the importance of continued monitoring and the viability of integrating processes affecting the productivity of natural systems with enhancement efforts. Finally, while this study does not evaluate fitness directly and lacks an unsupplemented control population, rates of genetic divergence measured here provide a range of multigenerational outcomes for contrasting management regimes. These comparative findings, in turn, can assist managers and policy-makers when assessing the relative benefits and risks of conservation decisions, particularly in cases where population recovery may depend on supportive breeding.

## Country-specific information (United States)

### Fisheries Genomics Work Funding Sources

NOAA – National Sea Grant Program, Saltonstall-Kennedy, National Science Foundation, Bonneville Power / Federal Columbia River Power System (FCRPS) Biological Opinion Remand Funds, US Department of Agriculture

### Funding sources accessed to support Genomics research

NOAA – National Sea Grant Program, Washington Sea Grant Program, National Science Foundation, Federal Columbia River Power System (FCRPS) Biological Opinion Remand Funds, US Department of Agriculture

### Examples of genetic/genomic information to inform fisheries management and/or policy decisions in USA

Many of the listings under the Endangered Species Act rely on genetic information. Such data has been used to delineate “Distinct Population Segments” (Conservation Units) that are the subject of management actions. There are many publications associated with this activity, including reports. NOAA maintains a list of technical reports on this website: https://www.nwfsc.noaa.gov/publications/scipubs/displayinclude.cfm?incfile=technicalmemorandum2016.inc

The publications of interest are the “status reviews”. The Hatchery Scientific Review Group on the West Coast of the USA has been extensively involved in developing “best practices” for the recovery and enhancement of salmon populations. http://www.hatcheryreform.us/hrp/welcomeshow.actionMany of the reforms are based on a wide number of papers that have been published on the impacts of hatchery fish on wild fish. Work in this area is also influencing the management of other fish species, as well as molluscan population management.

There is a growing interest in “parentage based tagging” (PBT) for tagging fish populations, which is viewed as an alternative for the coded wire tag program. PBT has significant potential to contribute to fisheries management.

Molecular-based methods are used extensively for stock identification, mixed stock analysis and measures of abundance

## Acknowledgement

We thank everyone who was involved in establishing CESRF, shaping its research direction, and sampling broodstock, including Levi George, Melvin Sampson, Steve Schroder, Craig Busack, past and present members of the Independent Scientific Review Panel, and the Yakama Nation Tribal Council. We are grateful to Maren Wellenreuther and Louis Bernatchez, and the OECD Co-operative Research Programme for the opportunity to present this research at the World Fisheries Congress in Korea. Funding for this study was provided by NOAA Fisheries/Federal Columbia River Power System (FCRPS) Biological Opinion Remand Funds (to K.A.N. and J.J.H.), Washington Sea Grant (Award NA14OAR4170078 to K.A.N), and the Hall Conservation Genetics Research Award from the University of Washington (to C.D.W.).

